# Identifying transcriptomic downstream targets of genes commonly mutated in Hereditary Hemorrhagic Telangiectasia

**DOI:** 10.1101/2022.11.25.517570

**Authors:** Md Khadem Ali, Yu Liu, Katharina Schimmel, Nicholas H. Juul, Courtney A. Stockman, Joseph C. Wu, Edda F. Spiekerkoetter

## Abstract

Hereditary Hemorrhagic Telangiectasia (HHT) is an autosomal dominant disease that causes arteriovenous vascular malformations (AVMs) in different organs, including the lung. Three genes, ENG (endoglin), ACVRL1 (ALK1) and SMAD4, all members of the TGF-β/BMPR2 signaling pathway, are responsible for over 85% of all HHT cases. However, how these loss-of-function gene mutations lead to AVMs formation and what common downstream signaling they target is unknown. Here, using a combination of siRNA-mediated gene silencing, whole transcriptomic RNA sequencing, bioinformatic analysis, transcriptomic-based drug discovery, endothelial cells functional assays and VEGF signaling analysis, and *ex vivo* precision cut lung slice (PCLS) cultures approach, we uncovered common downstream transcriptomic gene signatures of HHT-casing genes and identified promising drug for HHT. We found the commonly used BMPR2-signaling downstream target ID1 is not a common downstream target of all the three HHT genes knockdown in human pulmonary microvascular endothelial cells (PMVECs). We identified novel common downstream targets of all the three HHT-causing genes that were enriched for HHT-related biological process and signaling pathways. Among those downstream genes, LYVE1, GPNMB, and MC5R were strong downstream targets that could serve as a better common downstream target than ID1. Furthermore, using the common downstream upregulated genes (HHT disease signature) following HHT gene knockdown, we identified a small molecule drug, Brivanib, that reversed the HHT disease signature, and inhibited VEGF-induced ERK1/2 phosphorylation, proliferation, and angiogenesis in PMVECs and inhibited some of the upregulated HHT disease genes in PCLS. Our findings suggest that Brivanib could be an emerging new drug for HHT.

## Introduction

Hereditary hemorrhagic telangiectasia (HHT), also known as Rendu-Osler-Weber syndrome, is a complex genetic disease that causes abnormalities of blood vessel formation in the liver, lung, brain, skin, nasal mucosa, and gastrointestinal tract [1]. HHT is a rare disease, affecting approximately 1 in 5000-10000 people worldwide [2]. The key cardinal clinical manifestations of the disease include epistaxis (nose bleeding, present in 90% of cases), gastrointestinal hemorrhage, mucocutaneous telangiectasias (blood vessel dilations), and visceral arteriovenous malformations (AVMs). Clinically, HHT is generally diagnosed based on the four established Curacao Criteria; recurrent epistaxis, telangiectasia, family history, and visceral AVMs [3]. The diagnosis of HHT is confirmed when three or more criteria are met, while it is suspected if just two criteria are fulfilled. AVMs are large abnormal connections between arteries and veins bypassing the capillaries and result in severe bleeding when present in the nose, GI tract, and brain or cause paradoxical emboli and stroke when present in the lungs. AVMs in the lung (15-45%) and liver (>70%) are most common, while brain AVMs develop in 10-23% of HHT patients [4]. Currently, HHT therapies aim to lessen the disease’s symptoms. However, there is currently no mechanism-based targeted therapy available.

The majority (about >85%) of the HHT patients have loss of function mutations in either ENG (HHT type 1) or ACVRL1 (ALK1) (HHT type 2), but a small percentage of them (approximately <1%) have mutations in SMAD4 causing the combined juvenile polyposis/HHT syndrome. Interestingly, all three HHT-causing genes encode proteins belonging to the same TGFß superfamily of proteins. All three HHT gene products function in SMAD-dependent signaling pathways in which the activated Smad complex enters the cell nucleus and regulates transcriptional programs. Previous *in vivo* studies showed that any of the three HHT gene mutations could cause vascular malformations since knockout alleles for ENG, ACVRL1, and SMAD4 all result in HHT-like phenotypes in mice [1]. Mechanistically, in addition to the TGFß/BMP pathway, several other pathways, such as VEGF, mTOR, and PI3K/AKT signaling pathways, are associated with AVM formation and HHT pathogenesis [1]. Recently, pharmacologically combined treatment with Sirolimus and Nintedanib has been shown to correct Smad1/5/8 reduction and mTOR and VEGFR2 activation. This combination drug treatment reversed and prevented vascular abnormalities, bleeding and associated anemia in two experimental HHT mouse models (BMP9/10i antibody-, and the inducible ALK1 knock out mouse model of HHT) [5].

One of the known downstream targets of BMPR2/ALK1 signaling are the inhibitors of DNA-binding/differentiation proteins (IDs). ID proteins are thought to inhibit differentiation and promote cell cycle progression, to be involved in venous, arterial, and lymphatic endothelial cell identify and, therefore, when dysregulated, could possibly explain the aberrant proliferation observed during AVM pathogenesis in HHT[6]. In support of this, mice with a combined loss of both *Id1* and *Id3* display cranial hemorrhage secondary to the formation of an anastomosing network of dilated capillaries in the brain[7], suggesting that ID1/3 may play a role in AVM formation. Our group has previously identified the repurposed drugs Tacrolimus (FK506) and Enzastaurin as “ID1-increasing” drugs, which improved endothelial function *in vitro* [8, 9] as well as prevented and reversed the occlusive vasculopathy in pulmonary hypertension, a second rare disease characterized by haploinsufficiency in the BMPR2/ALK1 signaling pathway. FK506 was identified in a High Throughput Screen (HTS) of FDA-approved drugs using Id1 expression as a readout, whereas Enzastaurin was identified in a combined approach, using an siRNA screen (HTS) as well as *in silico* drug prediction of candidates that reverse the “transcriptomic disease signature” derived from blood cells from patients with pulmonary arterial hypertension (PAH). FK506 was used in an early proof of concept trial initiated at Stanford University [10] in stable PAH patients and improved outcomes when used as compassionate therapy in end-stage PAH patients [11].

Furthermore, low-dose FK506 attenuated nose bleeding in patients with HHT and PAH [12]. It was able to block the retinal pathology characterized by robust hypervascularization and AVM development in animal models of HHT[13]. While FK506 (and potentially Enzastaurin) might be beneficial in reducing AVM complications in HHT, we are proposing, that ID1 might not be the most specific readout for the dysfunctional ALK1/ENG/SMAD signaling in HHT, and therefore not the best target for drug repurposing efforts in HHT. The phenotypes of HHT (enlarged vascular malformations) and PAH (occlusive vasculopathy) are quite different, suggesting a different “gene expression signature” in HHT and PAH. Furthermore, given the complexity of the BMPR2/ALK1 pathway, including receptor heterodimerization, finely orchestrated ligand binding as well as interactions with other signaling pathways, mutations in BMPR2, as observed in PAH, likely do not result in the exact same signaling disturbances as mutations in ENG, ALK1 or SMAD4.

Since the three HHT-causing genes (ENG, ALK1, SMAD4) are responsible for the same phenotype, AVM formations, albeit with a different frequency and location in visceral organs depending on the specific mutation, it would be very important to identify downstream signaling abnormalities that are common to all three loss-of function mutations. The ultimate goal would be to identify ways to restore normal signaling and improve AVMs. Current therapies for large visceral AVMs in HHT are limited and consist of catheter-directed embolization or surgery, but no treatment exists that restores normal signaling. Drugs applied systemically that could restore normal signaling might prevent the development and growth of existing AVMs, an approach particularly important for children with HHT whose small AVMs grow with age. Furthermore, this approach might benefit patients with multiple smaller AVMs (in the lungs, nose, GI tract) and severe hypoxia, bleeding, and subsequent anemia, whose AVMs are not amenable to embolization because of their size and number but might respond to medical therapy, as has been shown with anti-angiogenic therapies including the use of VEGF inhibitors such as Bevacizumab for severe anemia[14]. Our study has two aims: ***First***, to identify common downstream genes and signaling pathways of the three known HHT-causing gene mutations (ALK1, ENG, SMAD4) using siRNA mediated gene downregulation, thereby mimicking complete local loss of function as seen in AVMs in HHT patients [15]. ***Second***, to identify repurposed/novel drugs that reverse the dysfunctional transcriptomic gene expression signature and normalize downstream signaling. We found that ID1 is not a common downstream target of all the three HHT gene mutations in PMVECs, and identified LYVE1, GPNMB, MC5R, and PLXDC2 as downstream targets of all three HHT genes. Importantly, we discovered a small molecule drug, Brivanib, that can activate downstream targets of ALK1/ENG/SMAD4 signaling, inhibit VEGF signaling pathways, and improve PMVECs functions.

## Methods

### Cell culture

Human healthy control pulmonary microvascular endothelial cells (PMVECs) (Cat # C12281, PromoCell GmbH, Heidelberg, Germany) were cultured in microvascular endothelial cell basal media (Cat # C-22120; PromoCell GmbH) supplemented with growth factors (Growth Medium MV SupplementPack, Cat # C-39220, PromoCell GmbH) and 100 U/mL Penicillin-Streptomycin Solution (Gibco) and used between passages 4 to 8. PMVECs were cultured under standard conditions (37°C, 5% CO_2_, 21% O_2_, 90% humidity).

### RNAi

Human pulmonary microvascular endothelial cells of passages 4-6 were seeded at 150K cells/well onto 6-well plates and incubated at 37ºC in a humidified 5% CO2 atmosphere. The next day, cells were washed with PBS and transfected with 50nM siRNAs against non-target controls, ACVRL1 (ALK1), ENG, SMAD4, LYVE1, GPNMB or MC5R (Thermo Fisher Scientific, Waltham, MA), and 2ul of Lipofectamine RNAiMAX in a total 1ml of OPTIMEM media. After 5 hrs of transfection, the medium was replaced with regular complete growth media. The following day, a starvation medium (0.2% FCS media) was added and incubated for 16 hrs. Cells were then stimulated with 20ng/ml of BMP9 for 2 or 24 hrs, harvested for RNA isolation, and performed RNAseq analysis.

### RNA sequencing (RNAseq)

RNA was isolated using RNeasy Plus Kits (Qiagen, Gaithersburg, MD) as per the manufacturer’s instructions. RNA samples were sent to the Novogene Corporation (Sacramento, CA), where the following steps were carried out: **Quality control:** Quality and integrity of total RNA were controlled on Agilent Technologies 2100 Bioanalyzer (Agilent Technologies; Waldbronn, Germany). The RIN values of all samples were in the ranges between 9.9-10. **Library construction:** The RNA sequencing library was constructed using NEBNext® Ultra II RNA Library Prep Kit (New England Biolabs) according to the manufacturer’s protocols. **Library quality control:** Library concentration was quantified using a Qubit 2.0 fluorometer (Life Technologies) and then diluted to 1ng/ul before checking insert size on an Agilent Technologies 2100 Bioanalyzer (Agilent Technologies; Waldbronn, Germany). The library was then quantified to greater accuracy by quantitative PCR (qPCR). **Sequencing:** 30 million paired reads for each sample were acquired with the Illumina NovaSeq 6000 system (Q20% (97-98%), Q30% (94-96%), GC content % (50-52%). **Data analysis**. The quality of the RNA-seq data was examined by base sequence quality plots using FastQC. TrimGalore was used to trim the sequence reads. Then, the RNA-seq reads were aligned to the human genome (hg19) using the STAR software, and a gene database was constructed from Genecode v19. Differentially expressed genes (DEG) between groups were quantified using the DESeq2 R package.

### Biological process, pathway enrichment, and protein-protein network analysis

Differentially expressed downstream genes of HHT knock-down (KD) conditions at 2 and 24h of BMP9 stimulations were uploaded on the ShinyGO 0.76 (http://bioinformatics.sdstate.edu/go/) platform to analyze GO biological process enrichment. The same data gene signatures were uploaded separately onto the DAVID Bioinformatics Resources 6.8 server (https://david.ncifcrf.gov/summary.jsp) for GO pathway enrichment analysis. The identifier was set to gene symbol, and *Homo sapiens* was selected to limit annotations in the gene list and background list. A significant value of *p* < 0.05 was set as the cutoff criterion. STRING Network analysis was performed using the SinyGO 0.76 platform.

### *In silico* Drug prediction

Following knockdown of the HHT-causing genes (ALK1, ENG, SMAD4) in PMVECs, we performed RNAseq and defined gene expression changes common to silencing of the three HHT-causing genes as “disease signature”. We then used the common downstream gene signature of all **up-**regulated genes 117 and 112 following HHT gene knockdown and at 2 and 24 h of BMP9 stimulation for drug prediction. Using the Broad Institute’s Clue.io query app https://clue.io/query, we identified compounds that reverse the common HHT disease signature of up-regulated genes. The list of up-regulated genes 117 (2h) and 112 (24h) genes following RNAseq was uploaded separately on the app https://clue.io/query, selecting with the gene expression L1000 query parameter. This web-based clue.io query app finds perturbagens that give rise to similar (or opposing) expression signatures. After running this query, we identified compounds that either mimic the transcriptional disease signature (compounds that could potentially worsen disease) or mimic the anti-signature (compounds that could potentially improve disease) in different cell lines. We only focused on findings from the human umbilical vein endothelial cells (HUVECs) data sets as this was the only dataset performed in endothelial cells, which are the critical cell type for AVM formation[16]. Top-ranked 5 CMAP compounds/drugs induced transcriptome alterations oppositional to (indicated by negative similarity mean) or overlapping with (indicated by positive similarity mean) caused by HHT causing gene knockdowns. Rank was determined by samples n>=3, tas value >=0.20, normalized connectivity score, and FDR value. Next, we narrowed down the top scoring drug list based on their relevance to VEGF inhibition as VEGF signaling is overactivated in HHT.

### Cell proliferation, apoptosis, and tube formation assay

MTT assay (Cat # V13154, Invitrogen, Waltham, MA), and caspase-3/7 assay (Cat # G8090, Promega, Madison, WI) were performed using commercially available kits as per the instructions to assess proliferation and apoptosis, respectively.

### Matrigel tube formation assay

The bottom of a 96-well plate was coated with 50ul per well of Matrigel and incubated at 37□°C for 1□h. PMVECs (1□×□10^4^ cells/100□μl per well) suspended in a starvation medium were added to the Matrigel and cultured at 37□°C and 5% CO_2_ for 5□h. Vessel tube-like structures were observed and photographed under a microscope with a digital imaging system. The data were analyzed using ImageJ with an angiogenesis analyzer tool.

### RNA isolation, RT-PCR, and qRT-PCR

Total RNA was isolated from cells using a commercially available RNeasy® Plus Mini Kit (Cat # 74134, Qiagen, Hilden, Germany).

For precision cut lung slices (PCLS), total RNA was extracted and purified using the TRizol RNA extraction protocol, as described previously [17]. The total RNAs were then converted to cDNA using a commercially available high-capacity cDNA Reverse Transcription Kit (Cat # 4368813, Applied Biosystems™, Foster City, CA) according to the manufacturer’s instructions. The expression levels of mRNAs were quantified using TaqMan™ 2x Universal PCR Master Mix (Cat # 4304437) and targeted Taqman probes (ALK1, ENG, SMAD4, GAPDH, Hs02786624_g1; ID1, Hs03676575_s1; HS3ST2, Hs00428644_m1; GPNMB, Hs01095679_m1; ANKRD33, Hs05002807_s1; RRAGD, Hs00222001_m1; LYVE1, Hs00272659_m1; CPA4, Hs01040939_g1; SLC25A47, Hs01584239_m1; SHISA9, Hs04188640_m1; HIST1H2BE, Hs00543841_s1; MC5R, Hs00271882_s1; FGF19, Hs00192780_m1; FRG2C, Hs01695863_sH; PLXDC2, Hs00262350_m1; KCNK5, Hs01123564_m1; and CACNA1G, Hs00367969_m1, Thermo Fisher Scientific, Waltham, MA) and normalized to a housekeeping control GAPDH.

### Western blot

Western blotting was carried out as described previously[8, 18]. The following antibodies were used: P-Smad1/5/9 (Cat # CST13820S, 1:1000), p44/42 MAPK (Erk1/2) (137F5) (ERK1/2 MAPK) (Cat # CST4695S, 1:1000), phosphorylated ERK1/2 (Phospho-p44/42 MAPK (Erk1/2) (Thr202/Tyr204) (E10), Cat # CST9106S, 1:1000), Id1 (sc133104, monoclonal, Santa Cruz Biotechnology, 1:100) and an HRP-conjugated secondary antibodies (Cat # ab205719 and ab6721, Abcam, 1:5000). The western blot band densitometric analysis was performed with ImageJ.

### Precision-cut lung slices (PCLS) culture

Lung tissue pieces were obtained from a healthy donor collected from the Donor Network West, San Ramon, CA. First, the tissue pieces were inflated with 2% low melting agarose prepared in 1x PBS, and kept on ice for 15mins. Then the solidified pieces were placed onto a petri dish plate and 8mm punch biopsies were carefully made, creating cylinders of lung tissue. 6% agarose was poured into the compress tome mold and the cylinder of lung tissue was quickly placed into the mold, ensuring that 6% agarose surrounded the cylinder on all sides. After being solidified, the tissue pieces were sliced using the compress tome at 400 μm thick slices. The lung slices were then cultured in 1x DMEM GlutaMAX medium containing 10% fetal calf serum, penicillin/streptomycin (1%), and amphotericin B (0.1%) in the presence and absence of Brivanib at 10 or 50 μM in 1mL of media in a 12-well tissue culture plate. After 24 hrs, the slices were washed with 1x PBS two times and then harvested for RNA isolation.

### Statistics analysis

All data analyses were performed using GraphPad Prism V9. All data are represented as the mean□±□standard error of the mean. An unpaired Student’s *t*-test was used to compare two groups, and one-way ANOVA was performed for comparing data with more than two groups, followed by an appropriate post-hoc test for multiple comparisons. Brivanib validation data in HHT gene knockdown cells were analysed using two-way repeated measures ANOVA with a Bonferroni post-hoc test. The statistically significant differences were considered at *p*□<□0.05.

## Results

### Identifying common downstream gene signatures of HHT-causing genes mutations in human PMVECs

As loss of function mutations in ALK1, ENG and SMAD4 are causative of HHT, and as all three genes belong to the TGFβ/BMPR2 super family signaling pathway (**Figure 1A**), we sought to identify common downstream signatures of all three HHT-causing gene mutations in PMVECs by RNAseq following siRNA-mediated silencing of the three genes. We used pulmonary microvascular endothelial cells, PMVECs, as endothelial cells are believed to be the critical cell type for AVM formation, and pulmonary cells, as pulmonary AVMs are one of the most common visceral manifestations in HHT. We first confirmed that following adding BMP ligand, BMP9, BMP signaling is activated in PMVECs, as evidenced by increased pSMAD1/5/9 and Id1 levels measured by western blot (**Figure 1B**). Next, we silenced ALK1, ENG, and SMAD4 with siRNA in PMVECs. As seen in **Figures 1C-E**, we achieved a knockdown of over 80% for all three genes, ALK1, ENG, and SMAD4, measured by qRT-PCR. We activated the pathway with 20ng/mL BMP9 for 2 and 24 hrs in the HHT genes knocked down cells. Stimulation with the ligand BMP9 had no effect on the expression of ALK1, ENG, and SMAD4 (**Figures 1C-E**). We then looked at one of the known downstream targets of the BMPR2/ALK1/ENG/SMAD4 pathway, Id1 (**Figure 1F and G**), and whether the expression of Id1 is reduced by knocking down all three HHT genes. Interestingly, while BMP9-induced Id1 expression is decreased by knocking down ALK1 and SMAD4, a knock-down of ENG did not affect BMP9-induced Id1 expression. In fact, Id1 expression was rather slightly increased after ENG knockdown compared to the NT siRNA condition. These findings indicated that Id1 is not a common downstream target of all three HHT genes but is specific for ALK1 and SMAD4 mediated signaling. This is a very important finding when it comes to identifying common downstream targets and subsequently predicting repurposed drugs that might increase the signaling for all three gene mutations.

**Figure 1.**
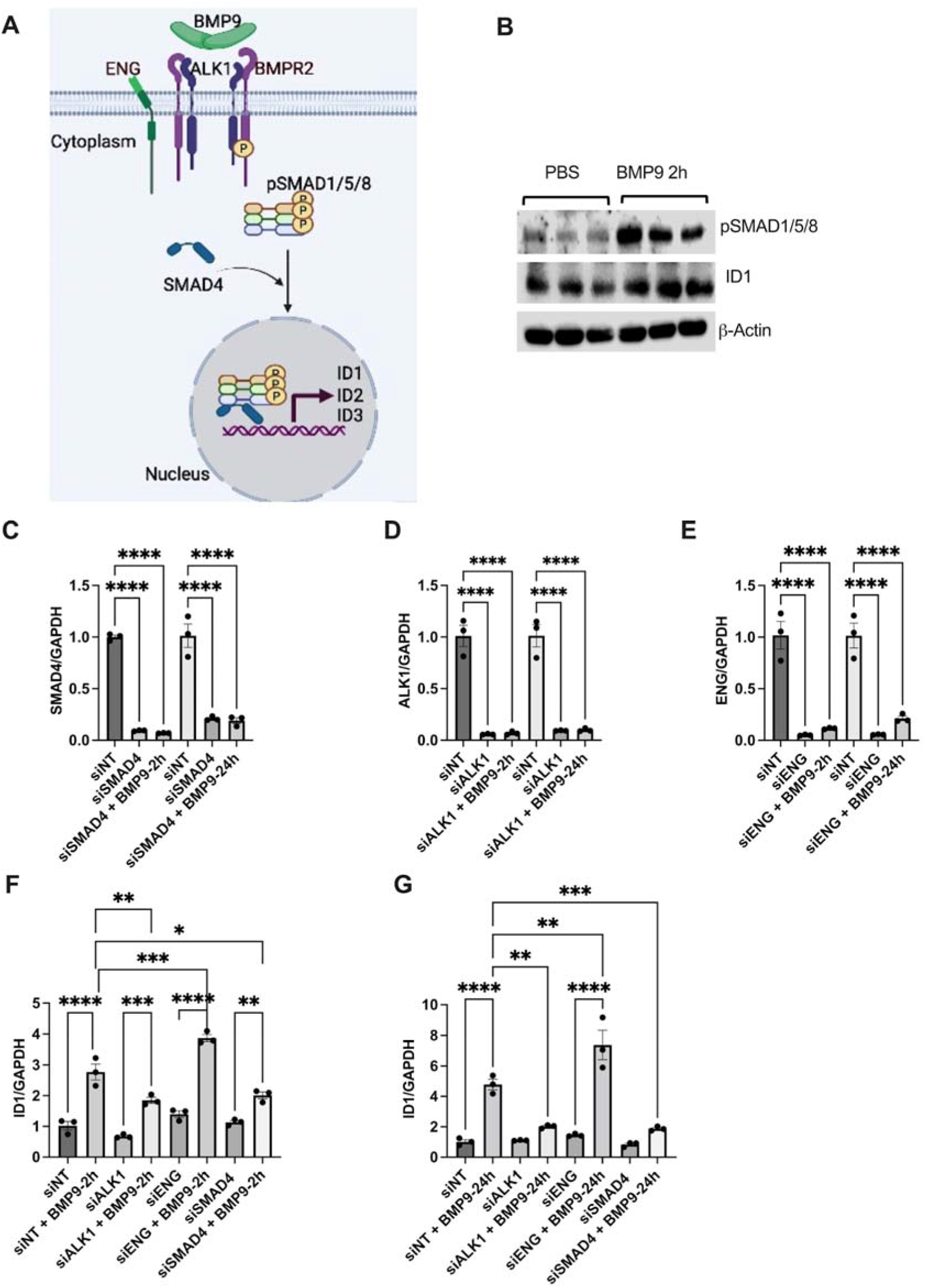
ID1 is not a common downstream target of ENG in PMVECs. A) Cartoon of BMP9 signaling in HHT. B) Western blot verification of BMP9 in PMVECs. C-E) siRNA-mediated knockdown verification of ALK1, ENG, and SMAD4 by qPCR. F and G) Validation of BMP9 signaling in ALK1, ENG, and SMAD4 knockdown conditions in PMVECs by qPCR (read out ID1).

Next, we carried out experiments to understand how knocking down ALK1, ENG, and SMAD4 effected the function of PMVECs *in vitro*. We assessed proliferation, apoptosis, and tube formation following the silencing of the three HHT genes with siRNA. The MTT assay showed that knockdown of the HHT genes did not significantly change cell proliferation (**Figure E1A**). Silencing of ENG induced apoptosis as evidenced by increased caspase 3/7 activity and decreased angiogenesis, as evidenced by reduced numbers of nodes, junctions and tube length compared to controls at baseline **(Figures E1B and C)**. ALK1 and SMAD4 knockdown had no significant effect on apoptosis and tube formation. These results suggested that while ALK1, ENG, and SMAD4 deficiency are linked with HHT, at baseline, only ENG deficiency induced PMVECs dysfunction *in vitro*.

Since HHT1 and HHT2 patients show similar clinical symptoms, a previous study had identified 277 downregulated and 62 upregulated common downstream targets genes in blood outgrowth endothelial cells isolated from HHT patients carrying ENG (HHT1), ALK1 missense or ALK1 non-sense mutations (HHT2) compared to healthy endothelial cells, which they called a “gene expression fingerprinting of HHT”[19]. However, to date, no studies have identified common downstream genes of the three key HHT genes in PMVECs. Thus, we performed RNA sequencing of PMVECs silenced to either ALK1, ENG, or SMAD4 and treated with BMP9 20ng/mL for 2 or 24 hrs. The list of significant differentially expressed genes was identified by employing the criteria of changes in gene expression of log2-fold changes value =>2 and *p* adjusted value =<0.05 between the groups. We identified 117 upregulated and 125 downregulated common genes 2hrs after BMP9 stimulation in PMVECs silenced for all three HHT genes (**Figures 2A and C**) and 112 upregulated and 132 downregulated common downstream genes 24 hrs after BMP9 stimulation in PMVECs silenced for all three HHT genes (**Figures 2B and C**). Consistent with previously identified genes associated with AVM/HHT[20, 21], we found a significant upregulation of ANGPT2 and APLN and downregulation of TMEM100 after 2hrs and 24hrs BMP9 stimulation in PMVECs knockdown cells (KD), respectively, along with other important vascular dysfunction related gene signature changes (**Figures 2A-C**).

**Figure 2.**
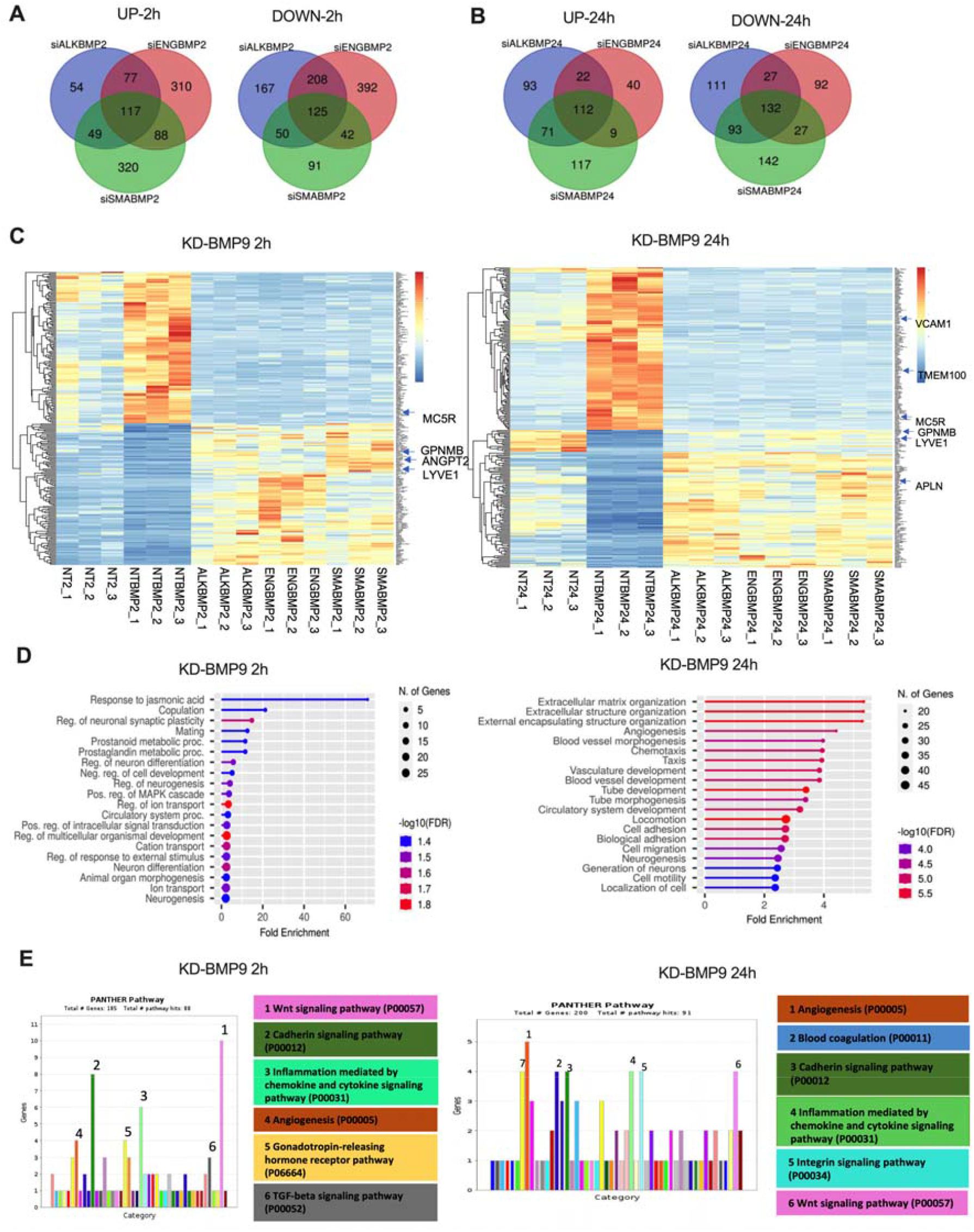
Common downstream gene signatures of ALK1, ENG and SMAD4 mutations enriched for AVM and HHT related biological processes and cell signaling pathways. **A and B)** Venn diagram of common upregulated or downregulated genes following knockdown of ALK1, ENG and SMAD4 in PMVECs stimulated with BMP9 (20ng/ml) for 2 or 24hrs (RNAseq). C) Heatmap of the common upregulated and downregulated genes following knockdown of ALK1, ENG and SMAD4 in PMVECs stimulated with BMP9 for 2 and 24hrs. D) SinyGo biological processes analysis of the common upregulated and downregulated genes following knockdown of ALK1, ENG and SMAD4 in PMVECs stimulated with BMP9 for 2 and 24hrs. E) Panther pathway analysis of the common upregulated and downregulated genes following knockdown of ALK1, ENG and SMAD4 in PMVECs stimulated BMP9 for 2 and 24hrs.

To determine whether the commonly identified downstream targets (“UP and Down genes”) of the HHT genes after 2hrs or 24hrs BMP9 stimulation could potentially associate with biological processes and signaling pathways relevant to HHT, we carried out an *in silico* analysis of the common downstream genes using the GO biological process enrichment analysis tool, the DAVID panther pathway analysis tool, and the STRING protein-protein interaction database [22]. GO enrichment analysis of the commonly dysregulated downstream targets uncovered a significant enrichment of HHT-related biological processes, including cell migration, adhesion, tube and vascular development and angiogenesis, extracellular matrix organization, and ion transport after 2 and 24hrs of BMP9 stimulation following HHT gene KD (**Figure 2D**). Moreover, we also observed enrichment of signaling pathways that are thought to regulate AVM formation in HHT, including Wnt signaling, cadherin signaling, integrin signaling, inflammation-mediated cytokines and chemokine signaling, blood coagulation, angiogenesis and TGFβ-signaling pathways (**Figure 2E**). STRING network analysis identified some hub genes, such as PDGFRB, and VCAM that could play roles in AVM/HHT (**Figures E2 A and B**).

Next, we identified the common persistently and consistently dysregulated gene signatures after 2 and 24 hrs of BMP9 stimulation in the common HHT gene KD conditions. We found 5 upregulated genes (Heparan Sulfate-Glucosamine 3-Sulfotransferase 2 (HS3ST2), Glycoprotein Nmb (GPNMB), Ankyrin Repeat Domain 33 (ANKRD33), Ras-related GTP-binding protein D (RRAGD), Lymphatic Vessel Endothelial Hyaluronan Receptor (LYVE1)) and 7 downregulated (Carboxypeptidase A4 (CPA4), Solute Carrier Family 25 Member 47 (SLC25A47), Shisa Family Member 9 (SHISA9), Histone cluster 1 H2B family member e (HIST1H2BE), Melanocortin 5 Receptor (MC5R), Fibroblast Growth Factor 19 (FGF19), FSHD Region Gene 2 Family Member C (FRG2C)) genes (**Figure 3A**). qRT-PCR validation of the common persistently dysregulated genes in PMVECs further confirmed upregulation of GPNMB and LYVE1 and downregulation of MC5R in the common HHT KD conditions after BMP9 stimulation. As further validation, those genes were regulated in the opposite direction when stimulated with BMP9 alone (**Figures 3B and C and Figures E3 A-D**). We therefore concentrated on MC5R, GPNMB, and LYVE1 to investigate further their known importance related to HHT. LYVE1 expression was positively correlated with preoperative edema in brain AVM [23], while the role of MC5R and GPNMB in AVM formation and HHT is unknown. Our in vitro functional assays did not exhibit significant changes in cell proliferation, apoptosis, and tube formation in PMVECs silenced to either MC5R, GPNMB or LYVE1 at baseline (**Figures E3 E-G**).

**Figure 3.**
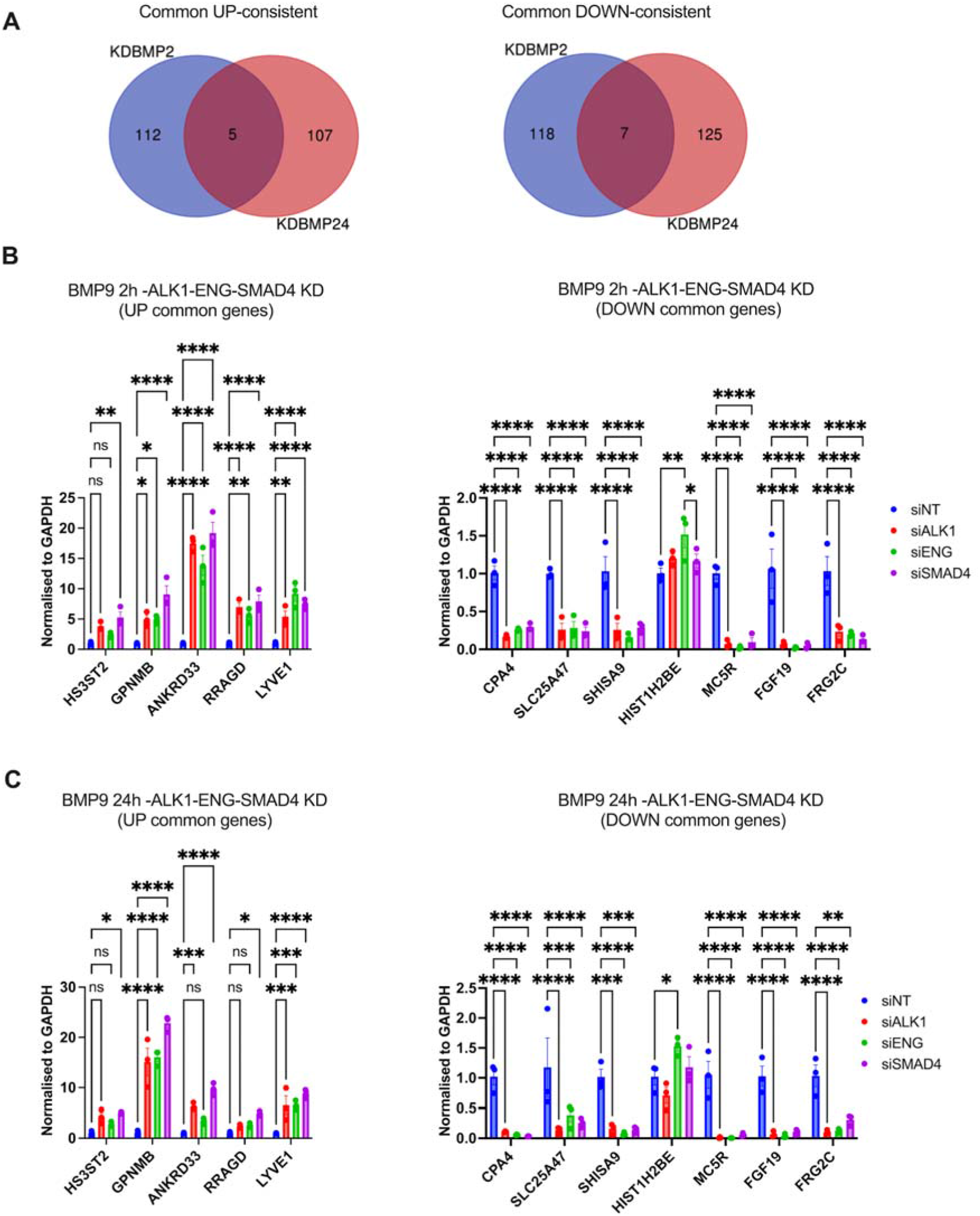
Identifying common persistently dysregulated downstream gene signatures of the HHT gene knockdowns in PMVECs. A) Venn diagram of the common persistent upregulated and downregulated downstream gene signatures between 2 and 24 hrs of BMP9 stimulation following HHT gene knockdown. B) qRT-PCR validation of the common persistent upregulated and downregulated downstream gene signatures at 2 hrs of BMP9 stimulation following HHT gene knockdown. C) qRT-PCR validation of the common persistent upregulated and downregulated downstream gene signatures at 24 hrs of BMP9 stimulation following HHT gene knockdown.

### In silico screening of drugs that can reverse common downstream gene signatures of HHT causing gene mutations

We hypothesized that novel/re-purposed drugs could reverse the HHT/PAVM pathological downstream targets/signaling and thereby reverse abnormal endothelial cell functions (tube formation, migration, proliferation) in PMVECs. A powerful strategy to predict novel drugs that might be beneficial in HHT is to use bioinformatics approaches to match in silico drug gene expression profiles with disease or anti-disease signature profiles, as previously shown by our group [8]. Here, we used the list of commonly upregulated genes that were differentially expressed in PMVEC silenced to all three HHT genes and stimulated with BMP9 for 2 and 24hrs, 117 and 112, respectively, which we labelled as “HHT disease signature”. This transcriptomic gene expression signature served as blueprint for predicting potentially beneficial drugs that mimic the complementary “anti-signature” using the Broad Institute’s Clue.io query app https://clue.io/query. We uploaded the genes on the web-based application database and selected with the gene expression L1000 query parameter. This web-based clue query app allows queries with external gene sets to identify compounds that either mimic our gene sets’ transcriptional signatures or reverse the signature (anti-signatures) in different cell lines. We narrowed down the drug list based on the findings from the HUVECs data sets as this is the closest cell line to our data sets acquired in PMVECs. The drugs were ranked following the criteria of samples >=3, tas values >=0.20, normalized connectivity score, and FDR values on the query app. Top scoring 5 HHT and anti-HHT drugs are shown in **Figure 4B**. From the top anti-HHT drug candidates (negative values) (24h), we concentrated on Brivanib as it is well-known for its anti-FGF/VEGF activities [24]. We hypothesized that Brivanib would on the one hand, reverse the dysfunctional downstream targets related to BMP signaling and on the other hand could inhibit the overactivated VEGF pathway in HHT. To test the effect Brivanib had on restoring members of dysfunctional BMP signaling, we determined whether Brivanib reversed the 5 upregulated (LYVE1, GPNMB, RRAGD, ANKRD33, HS3ST2) and 7 downregulated (MC5R, SLC25A47, FGF19, SHISA9, FRG2C, HIST1H2BE, CPA4) genes, we had previously defined a being consistently dysregulated after silencing of all three HHT genes and stimulation with BMP9 for 2 and 24hrs. Importantly, we observed that Brivanib inhibited the expression of LYVE1, GPNMB, RRAGD, and ANKRD33 induced in HHT gene KD conditions (**Figures 4C-F and E4 A-C**). Furthermore, Brivanib rescued the expression of the MC5R, SLC25A47, FGF19, SHISA9, and FRG2C that were downregulated in HHT gene KD conditions (**Figures 4G-K and E4 A-C**). These results demonstrated that Brivanib could improve the dysfunctional ALK1/ENG/SMAD4 signaling in HHT.

**Figure 4.**
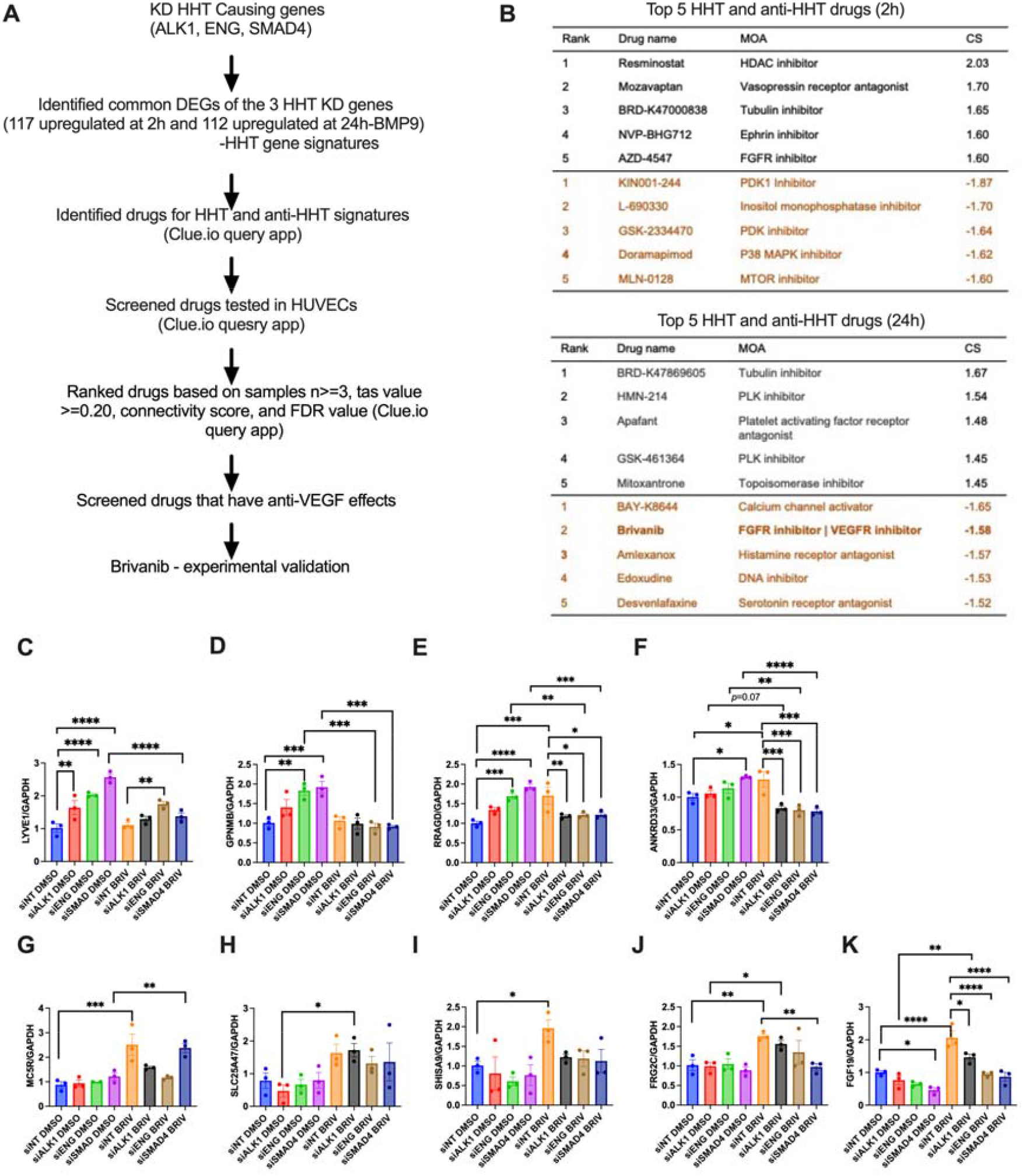
Drug prediction based on the common upregulated downstream targets after ALK1, ENG and SMAD4 knockdown and experimental validation in PMVECs. A) Experimental strategy: the common upregulated genes 117 (2h) and 112 (24h) after knockdown of the three HHT genes and stimulation with BMP9 for 2 and 24hrs (HHT disease signature) were uploaded separately on the Clue query app. The criteria of the drugs rank include samples >=3, normalized connectivity score, tas value >=0.20, and FDR value. B) top scoring 5 HHT (positive values) and anti-HHT drugs (negative values, indicated by brown color) are represented. (C-K) Effect of Brivanib on the expression of the common persistent downstream targets (LYVE1, GPNMB, RRAGD, ANKRD33, HS3ST2, MC5R, SLC25A47, FGF19, SHISA9, FRG2C, HIST1H2BE, and CPA4) after ALK1, ENG and SMAD4 knockdown was assessed by qRT-PCR in PMVECs. MOA, mode of action; cs, connectivity score; tas, transcriptional activity score. Data are represented as mean ± standard error mean (n=3). Two-way repeated measures ANOVA with a Bonferroni post-hoc test, \**P=*<0.5, \*\**P=*<0.01, \*\*\**P=*<0.001, \*\*\*\**P=*<0.0001.

### Brivanib inhibited the commonly upregulated downstream gene expression signatures of HHT-causing genes in ex vivo PCLS

To investigate whether Brivanib also inhibited the commonly upregulated signatures of HHT-causing genes in human lung tissue, we measured expression of LYVE1, GPNMB, HS3ST2, RRAGD, and ANKRD33 using qRT-PCR in human PCLS treated with Brivanib for 24 hrs. PCLS were prepared from a healthy human donor obtained from Donor Network West, California. PCLS were treated with 50 uM Brivanib or DMSO for 24 hrs and then harvested for RNA isolation (**Figure 5A**). Brivanib inhibited mRNA expression of LYVE1, GPNMB, and HS3ST2 in PCLS (**Figures 5B-F**). This inhibition in GPNMB, LYVE1and HS3ST2 expression *ex vivo* was similar to the inhibition of those genes in PMVECs after HHT gene knockout and Brivanib treatment *in vitro*. Brivanib did not change expression of RRAGD and ANKRD33 in PCLS.

**Figure 5.**
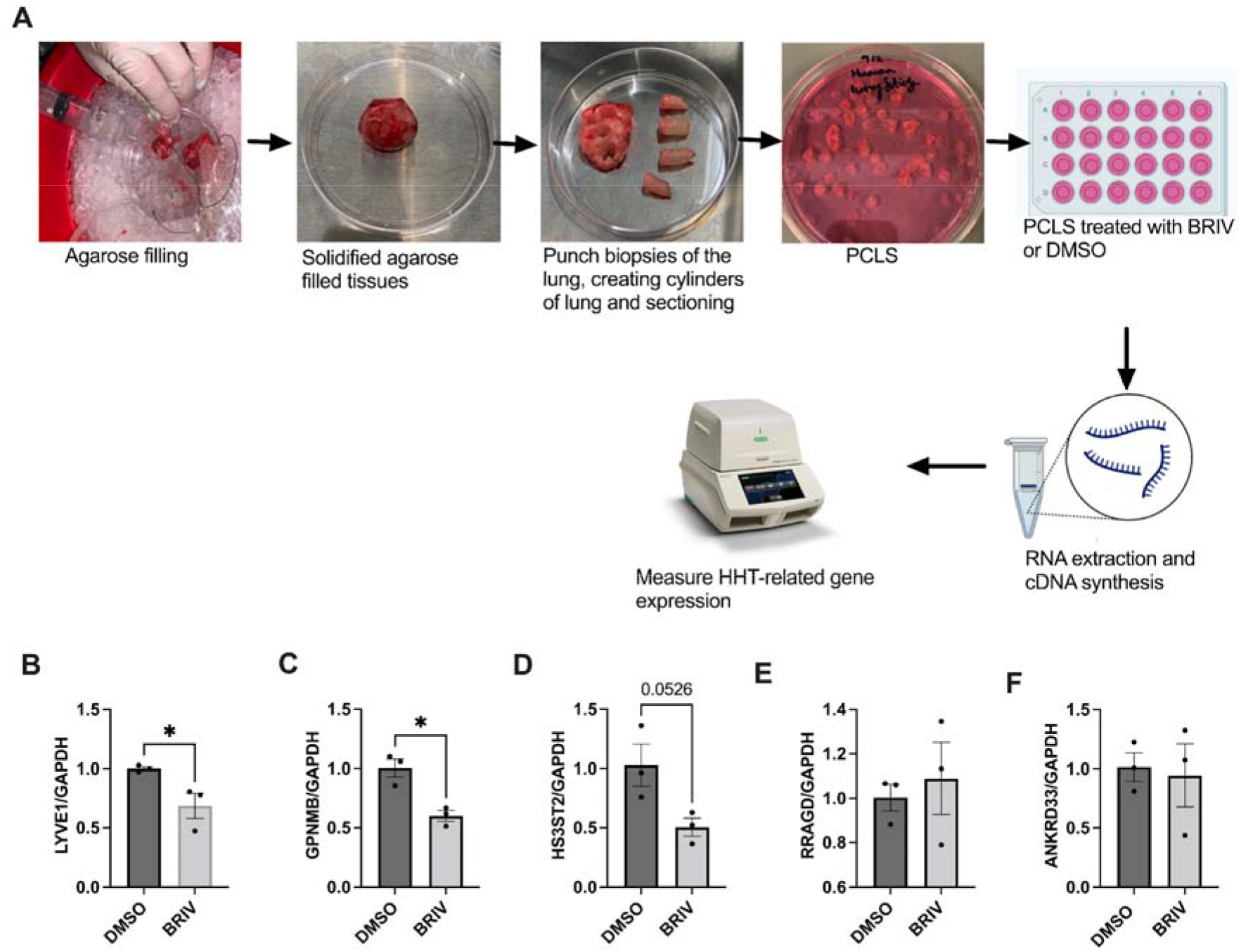
Effect of Brivanib on expression of upregulated disease signature genes in HHT in *ex vivo* healthy human PCLS. A) PCLS protocol. B) Expression of LYVE1, GPNMB, HS3ST2, RRAGD and ANKRD33 were measured by qRT-PCR following 24hrs treatment of Brivanib in PCLS (B-F). Data are represented as mean ± standard error mean (n=3). Unpaired Student’s *t*-test, \**P=*<0.5.

### Brivanib inhibited VEGF-induced ERK1/2 MAPK, -VEGF-induced PMVECs proliferation and tube formation *in vitro*

As over-activation of pro-angiogenic pathways, such as VEGF signaling, is strongly linked with the development of vascular dysfunction in HHT, and as Brivanib was previously shown to exhibit anti-VEGF effects both *in vitro* and *in vivo* in cancer and liver fibrosis studies [24-26], we carried out cell culture experiments to determine whether Brivanib can also inhibit the VEGF signaling pathway in PMVECs. PMVECs were cultured in the presence and absence of 10 μM Brivanib or DMSO in serum starvation media (0.2% FCS) for 24 hrs and then stimulated with 40 ng/mL VEGF for 10 mins. We measured phospho-ERK1/2 MAPK, total ERK1/2 MAPK (a downstream target of VEGF signaling) levels by western blotting. Notably, our results showed that Brivanib completely blocked VEGF-induced phospho-ERK1/2 levels in PMVECs (**Figure 6A**).

**Figure 6.**
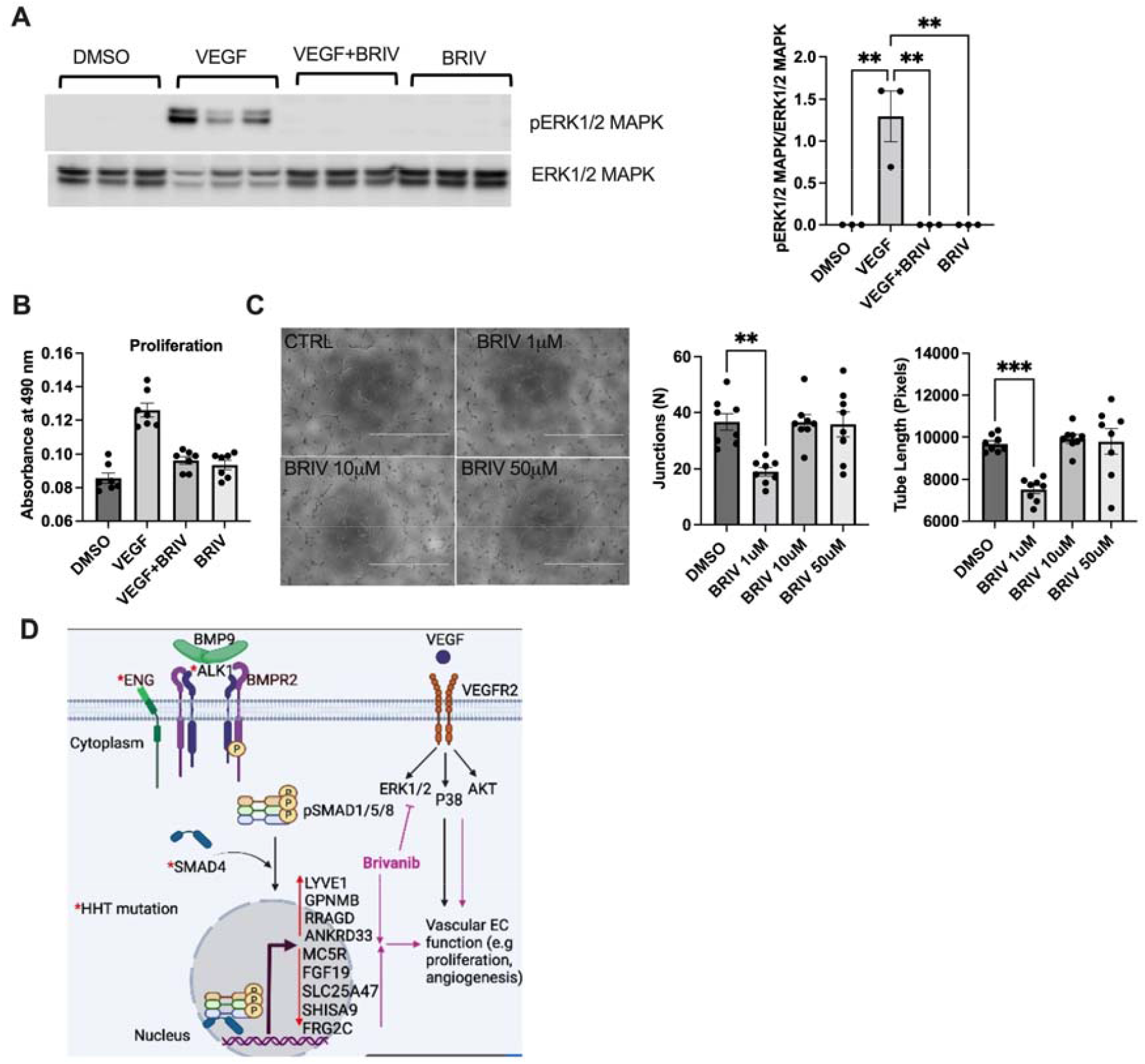
Brivanib inhibits VEGF-induced pERK1/2, -proliferation, and angiogenesis in PMVECs. A) PMVECs were treated with 10uM Brivanib or DMSO for 24 hrs followed by 20ng/mL VEGF or PBS for 10 mins. Protein was harvested and the effect of Brivanib on VEGF-induced phosphorylation of ERK1/2 was assessed in PMVECs by western blotting. B) The effect of Brivanib on VEGF-induced cell proliferation was assessed by MTT assay in PMVECs. C) Angiogenesis was assessed in a Matrigel tube formation assay using PMVECs treated with either Brivanib 1, 10, 50 uM or DMSO. D) Proposed model for the mechanism by which Brivanib might influence AVM and HHT pathogenesis.

We next assessed functional properties and endothelial behaviors with regards to tube formation and proliferation with and without Brivanib treatment. To determine whether Brivanib inhibits the VEGF-induced proliferation of PMVECs, we performed an MTT assay to measure the viability of PMVECs treated with VEGF (20 ng/mL) in the presence and absence of Brivanib (10 μM) in serum-starved media (0.2% FCS). As expected, VEGF treatment significantly increased PMVECs proliferation (**Figure 6B**). Importantly, Brivanib completely blocked the VEGF-induced proliferation of PMVECs (**Figure 6B**). Brivanib alone, without ligand stimulation, did not change proliferation of PMVECs. These findings suggested that Brivanib attenuated VEGF-induced PMVECs proliferation *in vitro*. To further explore the anti-angiogenic role of Brivanib on PMVECs *in vitro*, a matrigel based tube formation assay was performed. Results revealed that Brivanib inhibited tube formation in PMVECs as evidenced by the decreased number of junctions and total tube lengths compared to DMSO-treated control cells (**Figure 6C**).

## Discussion

In this study, we identified common downstream targets/pathways of the three HHT-causing genes, ALK1, ENG, and SMAD4 using whole genome RNAseq following knock down of the HHT genes in PMVECs *in vitro*. We also investigated whether we could identify a drug that can reverse the dysregulated downstream gene signatures and improve the common downstream dysfunctional pathways and HHT-related dysfunction of cellular phenotypes. Here, we found that ID1, a major downstream target of the TGFß/BMP signaling pathway, is not a common downstream target of all the HHT gene knockdown (ALK1, ENG, or SMAD4) conditions, while other downstream targets, such as LYVE1, GPNMB, PLXDC2, and MC5R, are downstream of all three HHT genes. Furthermore, we identified a small molecule drug, Brivanib, that could, on the one hand, reverse the downstream gene signatures following knockdown of all three HHT genes and, on the other hand, inhibit the VEGF signaling pathway, improve proliferation and tube formation in PMVECs. These findings suggest that Brivanib might be effective in treating AVMs and HHT by normalizing dysfunctional downstream signaling.

ID1 has been used as a downstream readout of BMPR2 signaling, and increasing ID1 as a therapeutic approach has been shown to be effective in PAH [8, 10]. HHT and PAH are both related rare genetic diseases, characterized by haploinsuffiency in members of the TGFß/BMPR2 signaling pathway. Mutations in BMPR2 cause PAH in up to 20% of patients, whereas mutations in ALK1, ENG and SMAD4 cause HHT in over 85% of patients.

Furthermore, there is an overlap between both diseases, as up to up to 10% of HHT patients show elevated pulmonary arterial pressures that indicate either the presence of Group 1 PAH (1% of HHT patients) or Group 2 PH associated with high output failure due to liver AVMs (more common, 10%) [27, 28].Our group previously discovered BMPR2 signaling inhibitors FHIT [8] and LCK [18], and BMPR2 signaling activating drugs FK506 [9] and Enzastaurin [8] by performing a high throughput screen (HTS) of siRNAs as well as FDA approved drugs to identify modifier genes and drugs that activate the BMPR2 pathway. Both screens were performed using the myoblastoma reporter cell line in which the BMP response element (BRE) from the ID1 promotor was linked to Luciferase (BRE-Luc). The BMPR2 signaling downstream target ID1 was the readout to measure activation of BMPR2 signaling in these HTS. As BMPR2 is the co-receptor of ALK1/ENG signaling we therefore first determined whether ID1 could be used as a common readout in HHT as well. We found that ID1 expression was not decreased by ENG knockdown, in contrast to ALK1 or SMAD4 knockdown, which inhibited BMP9 induced ID1 expression in PMVECs. While being a valid readout for ALK1 and SMAD4 mediated signaling, ID1 was not a readout for ENG signaling. In order to predict drugs that would normalize downstream signaling of all three HHT genes, we had to identify novel common downstream targets.

We used siRNA mediated, >80% knockdown of ENG, ALK1 or SMAD4 in PMVECs to mimic the proposed complete loss-of function of the above genes in endothelial cells of AVMs *in vivo*. While germline mutations in ALK1, ENG, or SMAD4 lead to haploinsufficiency and are required for AVM formation, several studies have suggested that they are not sufficient and that additional genetic and environmental factors are required to generate AVMs [29]. As an example, in order to establish skin and brain AVMs in adult mice lacking ALK1, the creation of a wound or stimulation with VEGF, two angiogenic triggers, were required for AVM formation [30-33]. It is proposed that these triggers might lead to a complete loss of function and signaling downstream of the HHT genes. This concept is further supported by the recent identification of somatic mutations in addition to germline mutations in endothelial cells of AVM lesions resulting in a bi-allelic loss of ENG or ALK1[15]. While the siRNA mediated complete loss of ENG, ALK1 or SMAD4 in PMVECs therefore mimics the *in vivo* loss of function, silencing of ENG, ALK1 or SMAD4 had no effect on PMVEC proliferation. Of interest ENG silencing decreased angiogenesis and induced apoptosis in PMVECs at baseline. Therefore the *in vitro* phenotype, without the use of additional stimuli such as VEGF (**Figure 6**) is not an adequate surrogate for the endothelial behavior after HHT KD in *vivo*.

To determine the common downstream targets of HHT-causing genes, we profiled gene expression by RNAseq following silencing of either ALK1, ENG, or SMAD4 in PMVECs. Our RNAseq analysis revealed, in addition to novel common downstream targets, several genes, such as ANGPT2, APLN, and TMEM100 (**Figure 2C**) that have previously been reported to be involved in AVM formation and HHT pathogenesis [20, 21]. This indicates that the list of common downstream genes could mimic AVM and HHT gene signatures. Angpt2 encodes for angiopoietin 2, an antagonistic ligand for TEK (TEK receptor tyrosine kinase, an EC surface receptor). Through RNAseq analysis Crist et al. found that Angpt2 was upregulated whereas Tek levels were downregulated in ECs isolated from the retina of *Smad4*-iECKO mice, a hereditary HHT mouse model. EC-specific Smad4 knockout resulted in an increase in Angpt2 transcription in those ECs that caused AVM formation in the retina [20]. Targeting Angpt2 with anti-ANGPT2 antibodies (LC-10) was shown to protect and rescue AVM formation of *Smad4*-iECKO mice, further supporting the role of ANGPT2 as a crucial TGFβ-downstream mediator of AVM development in the retina [20]. The same research team also found an upregulation of Apln (Apelin) in isolated lung ECs and retinas of *Smad4*-iECKO mice [34]. Apln is a ligand for the APJ receptor (also known as APLNR) which is a G-protein-coupled receptor. Previous studies showed a significant reduction of ECs proliferation and vascular outgrowth, abnormal arterial-venous alignment, and narrow blood vessels of mice and frog embryos deficient in either APLN or APLNR [35-41]. Hypoxia-induced APLN expression, and exogenous apelin treatment increased proliferation, migration, and inhibited apoptosis in mouse brain ECs *in vitro* [35]. APLN mRNA expression was also found to be downregulated by BMP signaling in human dermal microvascular ECs, and the BMP-APLN/APLNR signaling axis was crucial for hypoxia-induced ECs growth [42]. The APLN/APLNR pathway plays a significant role in various diseases, including pulmonary hypertension, and there is a strong relationship between BMP and APLN/APLNR in vascular signaling. However, the causal involvement of APLN/APLNR in AVM and HHT remains to be investigated. TMEM100 (encodes transmembrane protein 100) been shown to be enriched in arterial endothelium and activated by the BMP9/BMP10/ALK1 signaling axis [21, 43]. Intriguingly, mice lacking TMEM100 show substantial arterial specific abnormalities, embryonic lethality, and AVM formations, similar phenotypes seen in Alk1 constitutive KO mice [43, 44], suggesting that TMEM100 plays a significant role for arterial endothelium differentiation and vascular morphogenesis. Although Moon et al., claimed that TMEM100 is necessary for maintaining vascular integrity and angiogenesis, even though it is not the primary mechanism behind HHT pathogenesis, TMEM deficiency may contribute to the onset of HHT by weakening vascular integrity [21]. While there are no studies available that identify gene expression profiles of the HHT genes in PMVECs, three studies profiled genes expression in BOECs [19], HUVECs [45] and nasal telangiectasia tissue [46] of HHT patients. We compared our common gene list with the reported gene expression list of BOECs from HHT1 and HHT2 patients and found that 15 gene that were described to be dysregulated were common with our data set (PRCP, SLC40A1, MYO5C KCNH4, MGP, CPA4, MMP1, ANGPT2, ENG, APLN, HHIP, FABP4, AKR1C3, IGFBP3, and ESM1) [19]. Interestingly, among the 15 dysregulated genes, we only ENG and ESM1 (endothelial cell specific molecule 1) were changed in the same direction (upregulation) in the HHT BOECs when compared to our dataset. However MGP, MMP1, and TNFRS4 were upregulated and COL3A1 was downregulated in HUVECs of HHT1 and HHT2 patients as well as in our dataset [45]. The differences in altered gene expression profiles between our studies and the above-mentioned gene expression profiling studies could be explained by the fact that the authors used BOECs derived from HHT patients, which potentially were only haplo-insufficient for the specific gene mutation, whereas we used HHT gene-deficient PMVECs (80% knockdown). Together these findings suggest that the common downstream targets of the completely silenced HHT genes, we identified through RNAseq could potentially serve as a common HHT disease gene signature.

Furthermore, the top biological processes of the list of the common genes were significantly enriched for angiogenesis, blood vessels morphogenesis and development, vasculature development, tube morphogenesis and development, cell adhesion and migration, organ morphogenesis and development, regulation of signal transductions, and ECM organization (**Figure 2D**), all processes that are relevant to AVM formation and HHT pathogenesis [47]. The top signaling pathways enriched by the common genes list include Wnt, Cadherin, Integrin, TGFβ, chemokines and cytokines mediated inflammatory and angiogenesis pathways. While the involvement of TGFβ, inflammatory mediated signaling and angiogenic pathways in HHT is already known, the role of the Wnt signaling pathway in HHT still needs to be explored.

Next, by comparing two conditions, 2 and 24 hrs after BMP9 stimulation following silencing of the three HHT genes, we identified commonly altered gene signatures that were persistently dysregulated, LYVE1, GPNMB, MC5R and PLXDC2. We narrowed down the list of common persistent gene signatures by choosing those genes that showed an opposite direction of gene expression when stimulated with BMP9 for 2h and 24hrs versus when stimulated after complete gene knockdown. LYVE1 is expressed in lymphatic ECs but is also in developing blood vessels and macrophages. A previous study found increased expression of LYVE1 in human brain AVMs and the expression of LYVE1 was found to be positively associated with preoperative edema [23]. Glycoprotein non-metastatic melanoma protein B (GPNMB), a transmembrane protein, was shown to be linked with increased endothelial recruitment in a breast cancer study [48]. The activation of melanocortin receptors (MC1R and MC5R) in a mouse model of diabetic retinopathy improved retinal damage, prevented changes to the blood retinal barrier, and reduced local pro-inflammatory and pro-angiogenic factors, such as cytokines, chemokines, and VEGF [49]. PLXDC2, also known as tumor endothelial marker 7-related protein, TEM7R, is highly expressed in breast cancer and colon cancer tissues [50, 51]. These studies suggest that LYVE1, GPNMB, MC5R and PLXDC2 might play an important role in endothelial and vascular homeostasis in health and disease. Further studies are needed to confirm their causative role in AVM formation and HHT.

Currently, treatment options for HHT are limited. Several promising HHT drugs have been tested in clinical and preclinical settings. Bevacizumab (Avastin, anti-VEGF monoclonal antibody) [14, 52], Tacrolimus (FK506, a BMP signaling activator) [12, 53, 54], Pazopanib (TKI) [55], and Thalidomide (increases expression of PDGFB) [56] all have been shown to decrease epistaxis in HHT patients. In the genetically induced animal models of HHT, several drugs, such as Wortmannin, Pictilisib (a PI3K inhibitor) [57, 58], DC101 (an anti-VEGFR2 antibody)[59], SU5416 (a VEGFR2 inhibitor)[58], LC10 (a ANGPT2 inhibitor)[20], and G6.31 (a anti-VEGFA antibody) [32] have been shown to prevent or reduce retinal, peripheral or skin AVMs in neonate or adult mice. Recently, correcting multiple signaling pathways associated with AVM/HHT simultaneously, such as SMAD1/5/9, VEGFR2, and AKT/PI3K with combined treatment of drugs that target the above signaling (Sirolimus and Nintedanib) has also been shown to be effective in reducing and reversing retinal AVMs in the anti-BMP9/10 monoclonal antibody-induced HHT model in neonate and adult mice. While these drugs show promising findings in preclinical and clinical studies, several obstacles need to be addresses before a successful translation into the clinic can be made. As an example, FK506, an immunosuppressive drug, was demonstrated to improve vascular pathology in animal models [13], yet had minimal effects on epistaxis in a RCT HHT clinical trial when used as a topical ointment[53], while demonstration some efficacy yet also adverse effects when taken orally[54]. Further larger clinical studies are needed to confirm these findings. Furthermore, FK506 was discovered using ID1 as readout in the mouse myoblastoma BRE-Luc reporter cell line. As we found that ID1 is a common downstream target of ALK1 and SMAD4, but not necessarily ENG, this would suggest that FK506 may not be the optimal treatment for ENG-mutant HHT patients. In addition, while several clinical and animal studies document that anti-VEGF antibodies or VEGF inhibitors are effective to reduce bleeding events and anemia, the VEGF/VEGFR2 signaling pathway is complex and involves many downstream signaling pathway[60]. It is unclear which pathway (or combination of pathways) needs to be targeted precisely to facilitate regression of AVMs. Moreover, the understanding of the crosstalk between VEGF and BMP signaling is still limited, and how the two signaling pathways are disrupted in HHT is not completely known.

Our strategy to identify beneficial drugs for HHT is different from previous approaches. We predicted drugs based on their ability to reverse common downstream targets of HHT-causing genes and narrowed down the top candidates by introducing a second selection criterium: the drug should have in addition anti-VEGF properties. Using this approach, we selected three top-scoring potential HHT drugs (Brivanib, Cediranib, and Glesatinib). Among these drugs, we found that Brivanib can improve the dysfunctional common downstream gene signatures in PMVECs subjected to silencing of either ALK1, ENG or SMAD4 or in the *ex vivo* PCLS system. Brivanib also reversed VEGF-induced downstream signaling pathways and improved endothelial function *in vitro*. Although our findings all stem from *in vitro* or *ex vivo* studies, in the context of the drug discovery based on signaling pathways, our findings are comparable to the findings of a previously reported study in which the authors used a combined drug treatment to correct SMAD1/5/9, VEGF and mTOR signaling. We, on the other hand, identified Brivanib which was capable of correcting the downstream HHT as well as VEGF signaling simultaneously. Brivanib is a well-known VEGF signaling inhibitor [25, 61-63]. Our study suggests that Brivanib could be effective in HHT by correcting downstream targets of HHT causing gene signatures as well as VEGF signaling to positively influence AVM formation and growth (**Figure 6D**).

In summary, the present study used a RNAseq high throughput approach following loss of function mutations of the HHT causing genes to identify the common downstream gene signatures. Drugs were predicted based on their capability to mimic downstream HHT gene signatures as well as their anti-VEGF properties. Our findings suggest that ID1 is not a common downstream target of ENG but is specific for ALK1 and SMAD4 in PMVECs. We also revealed that Brivanib is superior to the use of a VEGF inhibition alone, as it restores normal HHT downstream signaling and thereby could be tested to prevent, halt or reverse AVMs. As all our findings of Brivanib are based on *in vitro* and *ex vivo* studies, Brivanib would need to be tested in *in vivo* HHT animal models and in clinical studies to further confirm these findings.

## Support statement

This research was supported by funding from the National Institutes of Health (R01 HL128734), Stanford Vera Moulton Wall Center for Pulmonary Vascular Diseases, and the U.S. Department of Defence (PR161256).

## Author contributions

Conceptualization: M.K.A., E.S.; Methodology: M.K.A; Data curation: M.K.A., Y.L., N.H.J., C.A.S., Z., M.; Writing - original draft: M.K.A.; Writing - review & editing: M.K.A., E.S., K.S.; Supervision: E.S., Funding acquisition: E.S.

## Acknowledgement

The authors are thankful to Dr. Adam M. Andruska, Division of Pulmonary, Allergy and Critical Care Medicine, Stanford University, for his critical comments and constructive suggestions to improve this manuscript.

## Conflict of Interest statement

The authors declare no competing or financial interests.

A patent application for the use of Brivanib in HHT is currently filed by Stanford University.

## Figure legends

**Figure E1.**
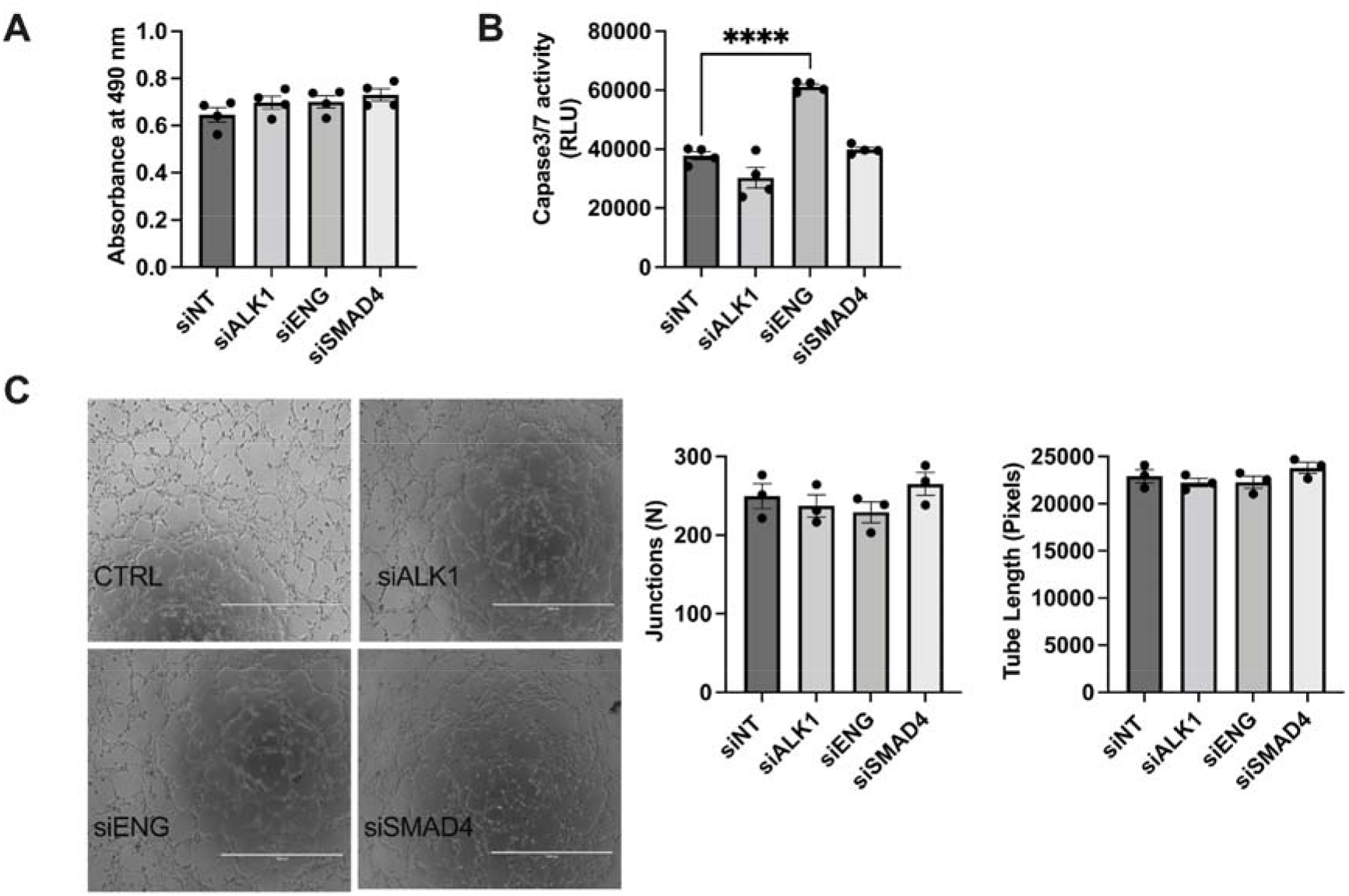
Functional consequence of ALK1, ENG, and SMAD4 knockdown in PMVECs (proliferation, apoptosis and tube formation).

**Figure E2.**
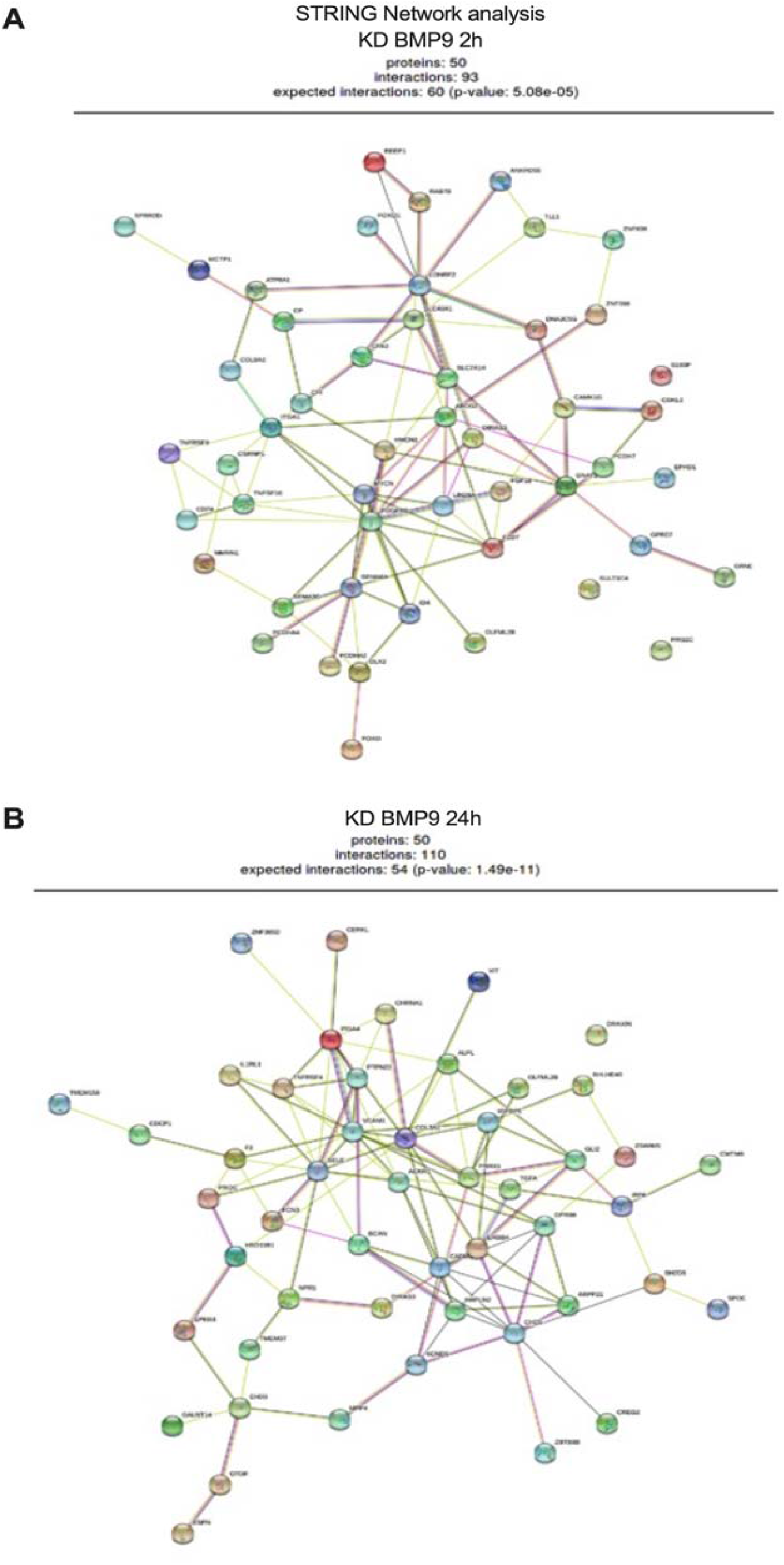
STRING network analysis of the common upregulated and downregulated genes following knockdown of ALK1, ENG and SMAD4 in PMVECs stimulated with 2h and 24h of BMP9.

**Figure E3.**
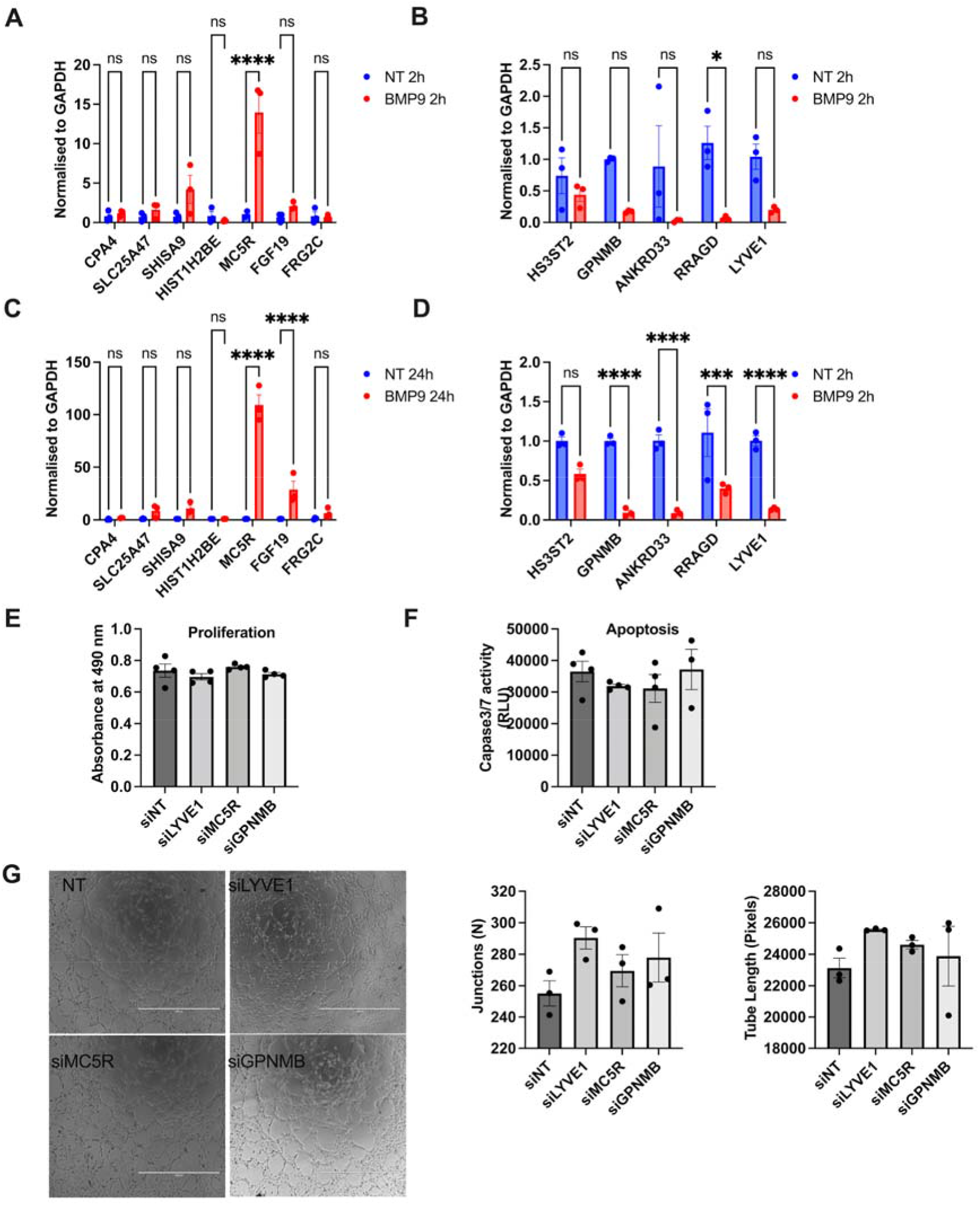
Validation of common upregulated or downregulated genes signatures with BMP9 stimulation at 2 (A and B) or 24 hrs (C and D) and functional consequences of the knockdown of LYVE1, GPNMB and MC5R in PMVECs. After 72 hrs knockdown of LYVE1, MC5R, and GPNMB PMVECs proliferation (E), apoptosis (F), tube formation (G) were assessed.

**Figure E4.**
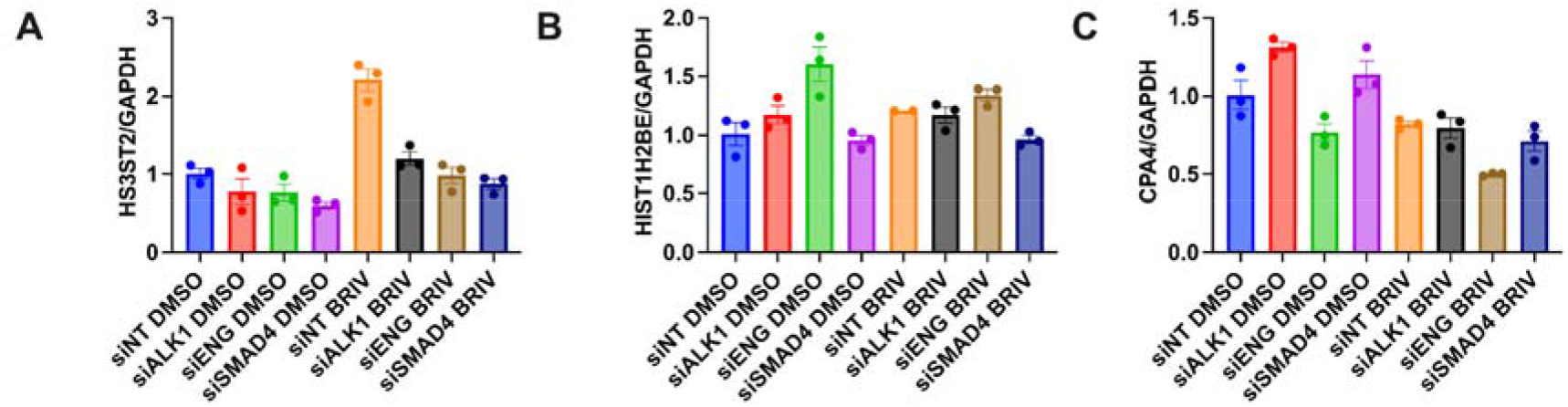
Effect of Brivanib on the expression of the common persistent downstream targets after ALK1, ENG and SMAD4 knockdown were assessed by qRT-PCR in PMVECs (A-C).

